# Homozygous loss of autism-risk gene *CNTNAP2* results in reduced local and long-range prefrontal functional connectivity

**DOI:** 10.1101/060335

**Authors:** Adam Liska, Alice Bertero, Ryszard Gomolka, Mara Sabbioni, Alberto Galbusera, Noemi Barsotti, Stefano Panzeri, Maria Luisa Scattoni, Massimo Pasqualetti, Alessandro Gozzi

## Abstract

Functional connectivity aberrancies, as measured with resting-state fMRI (rsfMRI), have been consistently observed in the brain of autism spectrum disorders (ASD) patients. However, the genetic and neurobiological underpinnings of these findings remain unclear. Homozygous mutations in Contactin Associated Protein-like 2 (CNTNAP2), a neurexin-related cell-adhesion protein, are strongly linked to autism and epilepsy. Here we used rsfMRI to show that homozygous mice lacking *Cntnap2* exhibit reduced long-range and local functional connectivity in prefrontal and midline brain “connectivity hubs”. Long-range rsfMRI connectivity impairments affected heteromodal cortical regions and were prominent between fronto-posterior components of the mouse default-mode network (DMN), an effect that was associated with reduced social investigation, a core “autism trait” in mice. Notably, viral tracing revealed reduced frequency of prefrontal-projecting neural clusters in the cingulate cortex of *Cntnap2*^−/−^ mutants, suggesting a possible contribution of defective mesoscale axonal wiring to the observed functional impairments. Macroscale cortico-cortical white matter organization appeared to be otherwise preserved in these animals. These findings reveal a key contribution of ASD-associated gene CNTNAP2 in modulating macroscale functional connectivity, and suggest that homozygous loss-of-function mutations in this gene may predispose to neurodevelopmental disorders and autism through a selective dysregulation of connectivity in integrative prefrontal areas.

## Introduction

Neuroimaging and post-mortem studies have consistently revealed impaired or atypical connectivity across brain regions of autistic spectrum disorders (ASD) patients (Anagnostou and Taylor, 2011). These findings have led to the hypothesis that aberrant connectivity patterns might represent a common final pathway or neurobiological pathogenetic correlate of the autistic phenotype to which different ASD etiologies may converge (Just et al., 2012). Although great heterogeneity exists in the sign and distribution of abnormal connectivity across studies and imaging modalities, consistent features indeed appear to emerge, including reduced functional coherence of long-range intra-hemispheric cortico-cortical default mode circuitry, impaired inter-hemispheric regulation and possible increase in local and short-range cortico-subcortical coherence (Rane et al., 2015). However, the neurophysiological underpinnings of these connectional derangements are largely unknown, and a causal etiopathological contribution of specific genetic variants to impaired connectivity in ASD remains to be firmly established.

Mouse lines recapitulating high-confidence ASD mutations (Sanders et al., 2015) have been employed to understand how specific genetic alterations translate into relevant changes in cells and circuits (Auerbach et al., 2011). The recent optimization of neuroimaging readouts of functional connectivity such as resting-state functional MRI (rsfMRI) in the mouse (Sforazzini et al., 2014a) permits to extend this paradigm to the investigation of the elusive genetic and neurobiological foundations of aberrant connectivity observed in ASD (Liska and Gozzi, 2016). The approach leverages on the identification of robust homotopic and distributed rsfMRI connectivity networks in the mouse, including possible homologues of distributed human rsfMRI systems like the salience and default mode (DMN) networks (Gozzi and Schwarz, 2015), and the observation that cyto-architecturally conserved heteromodal cortices in cingulate and retrosplenial regions exhibit similar “hub-like” topological properties in both species (Cole et al., 2010; Liska et al., 2015). Importantly, as mouse rsfMRI measurements rest on the same biophysical principles as corresponding human neuroimaging readouts, this approach has the merit of providing a direct translational bridge across species.

Homozygous loss-of-function mutations in *Contactin Associated Protein-like 2 (CNTNAP2)*encoding CASPR2, a neurexin-related cell-adhesion molecule, are strongly linked to autism and epilepsy in consanguineous families (Strauss et al., 2006; Alarcin et al., 2008; Rodenas-Cuadrado et al., 2014). Loss of *Cntnap2* in mice leads to abnormal neuronal migration, reduced GABAergic neurons, spontaneous seizures, and behavioural traits consistent with ASD symptoms in humans (Penagarikano et al., 2011), an ensemble of traits that phenocopy major neuropathological features observed in cortical dysplasia-focal epilepsy (CDFE) syndrome, a rare neuronal migration disorder associated with a recessive mutation in *CNTNAP2* (Strauss et al., 2006). Interestingly, common genetic variants in *CNTNAP2* were recently described to be associated with impaired frontal lobe connectivity in humans (Scott-Van Zeeland et al., 2010). However, a causal relationship between ASD-related loss-of-function mutations in *CNTNAP2* and functional connectivity remains to be firmly established. Moreover, the role of *CNTNAP2* in shaping macroscale circuit assembly, and the specific substrates affected, remain largely unknown.

To address these questions, we used BOLD rsfMRI, diffusion-weighted MRI and retrograde viral tracing to map large-scale functional connectivity and white matter topology in homozygous *Cntnap2-null* mice (*Cntnap2^−/−^*). We document that loss of *Cntnap2* results in local and long-range connectivity reductions affecting prefrontal regions that act as “functional connectivity hubs” in the mouse brain (Liska et al., 2015), and that fronto-posterior *hypo-connectivity* is associated with impaired social behaviour. The presence reduced prefrontal-projecting neuronal frequency in the cingulate cortex of *Cntnap2^−/−^* mutants suggest a possible contribution of defective *mesoscale* axonal wiring to the observed functional connectivity impairments. Collectively, these results reveal a role of autism-risk gene *CNTNAP2* in modulating functional network assembly between key integrative connectivity hubs of the mammalian brain. The observed long-range prefrontal hypo-connectivity in *Cntnap2^−/−^* mice recapitulates imaging findings in autism and adds to the construct validity of this mouse line to model ASD-related phenotypes.

### Materials and methods

#### Ethical statement

All *in vivo* studies were conducted in accordance with the Italian law (DL 116, 1992 Ministero della Sanità, Roma) and the recommendations in the Guide for the Care and Use of Laboratory Animals of the National Institutes of Health. Animal research protocols were also reviewed and consented to by the animal care committee of the Istituto Italiano di Tecnologia. The Italian Ministry of Health specifically approved the protocol of this study, authorization n° 07753 to A.G. All surgical procedures were performed under anaesthesia.

#### Animals

*Cntnap2*-null (*Cntnap2^−/−^*) and control “wild-type” (*Cntnap2^+/+^*) breeding pairs were obtained from Jackson Laboratories (Bar Harbor, ME, USA) and bred locally. Mice were housed by sex in mixed genotype groups, with temperature maintained at 21 ± 1°C and humidity at 60 ± 10%. All experiments were performed on adult male mice between 12-16 week of age, corresponding to young maturity. The specific age-range for each experimental activity is reported below. No onset of spontaneous seizures was observed in any of the *Cntnap2* mutants or control mice tested in behavioural, imaging or tracing studies. This is consistent with previous reports showing propensity for spontaneous epileptic episodes in *Cntnap2^−/−^* to occur only after 6 months of age (Penagarikano et al., 2011).

#### Social interaction

For behavioural testing, 12-week-old *Cntnap2^−/−^* and control *Cntnap2^+/+^* mice (n = 13 each group), were evaluated in the male-female social interaction test during the light phase, as previously described (Scattoni et al., 2011; Scattoni et al., 2013). An unfamiliar stimulus control female mouse in estrous was placed into the home-cage of an isolated test male mouse, and social behaviour were recorded during a 3-min test session. Scoring of social investigation parameters was conducted using Noldus Observer 10XT software (Noldus Information Technology, Leesburg, VA, USA). Social interactions were defined as number of events (frequency) and duration of the following behavioural responses performed by the test mouse: anogenital sniffing (direct contact with the anogenital area), body sniffing (sniffing or snout contact with the flank area), head sniffing (sniffing or snout contact with the head/neck/mouth area), locomotor activity, rearing up against the wall of the home-cage, digging in the bedding, and grooming (self-cleaning, licking any part of its own body). Social investigation is defined as the sum of sniffing and following behaviours (Scattoni et al., 2008). No observations of mounting, fighting, tail rattling, and wrestling behaviours were observed. Scoring was rated by two investigators blind to genotype. Inter-rater reliability was 98%. To measure ultrasound vocalization recordings, an ultrasonic microphone (Avisoft UltraSoundGate condenser microphone capsule CM16, Avisoft Bioacoustics, Berlin, Germany) was mounted 20 cm above the cage and the USVs recorded using Avisoft RECORDER software version 3.2. Settings included sampling rate at 250 kHz; format 16 bit. The ultrasonic microphone was sensitive to frequencies between 10 and 180 kHz. For acoustical analysis, recordings were transferred to Avisoft SASLabPro (version 4.40) and a fast Fourier transformation (FFT) was conducted as previously described (Scattoni et al., 2008). Start times for the video and audio files were synchronized.

#### Resting-state fMRI

rsfMRI experiments were performed on the same experimental cohorts employed in the behavioural tests (*n* = 13 Cntnap2^+/+^; *n* = 13 Cntnap2^−/−^). At the time of imaging, mice were 13-14 weeks old. The animal preparation protocol was recently described in great detail (Ferrari et al., 2012; Sforazzini et al., 2014b). Briefly, mice were anaesthetized with isoflurane (5% induction), intubated and artificially ventilated (2% maintenance). The left femoral artery was cannulated for continuous blood pressure monitoring and terminal arterial blood sampling. At the end of surgery, isoflurane was discontinued and substituted with halothane (0.75%). Functional data acquisition commenced 45 min after isoflurane cessation. Mean arterial blood pressure was recorded throughout imaging sessions. Arterial blood gases (p_a_CO_2_ and p_a_O_2_) were measured at the end of the functional time series to exclude non-physiological conditions. Mean p_a_CO_2_ and p_a_O_2_ levels recorded were 17 ± 3 and 250 ± 29 mmHg in *Cntnap2^+/+^* and 15 ± 3 and 231 ± 38 mmHg in *Cntnap2^−/−^*. Possible genotype-dependent differences in anaesthesia sensitivity were evaluated using Student’s two-sample t-test applied to two independent readouts previously shown to be linearly correlated with anaesthesia depth: mean arterial blood pressure and amplitude of cortical BOLD signal fluctuations (Steffey et al., 2003; Liu et al., 2011; Zhan et al., 2014).

rsfMRI images were acquired with a 7.0 Tesla MRI scanner (Bruker Biospin, Milan) as previously described (Liska et al., 2015), using a 72 mm birdcage transmit coil and a four-channel solenoid coil for signal reception. For each session, high-resolution anatomical images were acquired with a fast spin echo sequence (repetition time (TR)/echo time (TE) 5500/60 ms, matrix 192 × 192, field of view 2 × 2 cm^2^, 24 coronal slices, slice thickness 0.50 mm). Co-centred single-shot BOLD rsfMRI time series were acquired using an echo planar imaging sequence with the following parameters: TR/TE 1200/15 ms, flip angle 30°, matrix 100 × 100, field of view 2 × 2 cm^2^, 24 coronal slices, slice thickness 0.50 mm, 500 volumes and a total rsfMRI acquisition time of 10 min. Readers can contact the corresponding author for access to the MRI raw data, templates and code employed to generate the functional maps.

#### Functional connectivity analyses

The first 20 volumes of the rsfMRI data were removed to allow for T1 equilibration effects. The time series were then despiked, corrected for motion and spatially normalized to an in-house mouse brain template (Sforazzini et al., 2014a). The normalised data had a spatial resolution of 0.1042 × 0.1042 × 0.5 mm^3^ (192 × 192 × 24 matrix). Head motion traces and mean ventricular signal (averaged rsfMRI time course within a reference ventricular mask) were regressed out of each of the time series. No inter-group differences in ventricular volume was observed as measured by the dimension of individual ventricular masks (t-test, p = 0.31). All rsfMRI time series were spatially smoothed (full width at half maximum of 0.6 mm) and band-pass filtered to a frequency window of 0.01-0.1 Hz.

To obtain an unbiased identification of the brain regions exhibiting genotype-dependent differences in functional connectivity, we implemented recently developed aggregative metrics for these parameters (Cole et al., 2010; Maximo et al., 2013; Liska et al., 2015) and calculated local and global connectivity maps for all subjects. This metric considers connectivity of a given voxel to a subset of all other voxels within the brain mask by computing average connectivity strength. Specifically, we employed the weighted connectivity method, in which individual *r*-scores are first transformed to *z*-scores using Fisher’s *r*-to-*z* transform and then averaged to yield the final connectivity score. Local connectivity strength was mapped by limiting this measurement to connections within a 6-voxel radius sphere (0.6252 mm in plane), while long-range connectivity was computed by considering only connections to voxels outside this sphere. The radius employed represents approximately half the thickness of mouse anterior cortex (Dodero et al., 2013) and is a good approximation of the overall average cortical thickness (Braitenberg and Schüz, 2013; Sun and Hevner, 2014). The use of this value ensures that the employed local connectivity metric reflects purely intra-cortical effects at least in outmost cortical voxels and in thicker fronto-cortical regions. This value is proportionally much lower than what is commonly employed in human local connectivity mappings, where values as large as 14 mm (i.e., 4/5-fold mean human cortical thickness) have been employed (reviewed by Maximo et al., 2013).

Voxelwise inter-group differences in each of these parameters were mapped using a two-tailed Student’s *t*-test (p < 0.05 FWE cluster corrected with cluster-defining threshold of *t*_24_ > 2.06, p < 0.05, as implemented in FSL). The effect was also quantified in volumes of interest (VOIs). The anatomical location of the examined VOIs is reported in Figure S1. Region identification and naming follow classic neuroanatomical labelling described in (Paxinos and Franklin, 2011). Many of these regions have recently been reclassified according to their cytoarchitectural properties such to match analogous regions in human and primates (Vogt and Paxinos, 2014oyed (reviewed by Maxim). According to this scheme, the mouse prelimbic cortex corresponds to Brodmann area 32 (A32), cingulate cortex area 1 (anterior cingulate cortex) to Brodmann area A24b, infralimbic cortex to A24a, retrosplenial cortex to areas A30 and A29. In keeping with this and the comparative work of other authors (Öngür and Price, 2000), in this paper we define the mouse prefrontal cortex (PFC) as an anatomical ensable of regions inluding prelimbic, infralimbic and anterior cingulate cortex, corresponding to Brodmann areas A24a/b, A32, and A10.

Inter-group differences in the extension and intensity of long-range rsfMRI correlation networks were mapped using seed-based approach as previously described (Sforazzini et al., 2014b). Small *a priori* seed regions of 3×3×1 voxels were chosen to cover antero-posterior cortical networks and representative heteromodal cortical structures (Fig. S2). The mean time courses from the unilateral (medial, Rs, PrL) and bilateral seeds (TeA, Pt, and vHC) were used as regressors for each voxel. Group level differences in connectivity distributions were mapped using two-tailed Student’s t-tests (p < 0.05 FWE cluster corrected with cluster-defining threshold of *t*_24_ > 2.06, p < 0.05, as implemented in FSL).

Alterations in inter-hemispheric functional connectivity were assessed by computing correlation coefficients of inter-hemispheric VOI pairs depicted in Fig. S1. The statistical significance of inter-group correlation strength in each VOI was assessed with a two-tailed Student’s *t*-test (*t*_24_ > 2.06, p < 0.05) and corrected for multiple comparisons using a false discovery rate q = 0.05 according to the Benjamini-Hochberg procedure.

Anteroposterior DMN connectivity was mapped by computing seed-to-VOI correlations. Prelimbic and cingulate cortex were employed as prefrontal volumes of interest. The location of seeds employed for mapping are indicated in Fig. S2. The statistical significance of inter-group effects was quantified using a two-way repeated-measures ANOVA, where seed location and genotype were used as variables.

#### Diffusion MRI

*Ex vivo* diffusion-weighted (DW) MRI was carried out on paraformaldehyde fixed specimens as previously described (Dodero et al., 2013). At the end of the rsfMRI experiments, mice were transcardially perfused with 4% para-formaldehyde under deep isoflurane anaesthesia. Brains were imaged inside intact skulls to avoid post-extraction deformations. Each DW dataset was composed of 8 non-diffusion-weighted images and 81 different diffusion gradient-encoding directions with b=3000 s/mm^2^ (δ=6 ms, Δ=13 ms) acquired using an EPI sequence with the following parameters: TR/TE=13500/27.6 ms, field of view 1.68 × 1.54 cm^2^, matrix 120 × 110, inplane spatial resolution 140 × 140 μm^2^, 54 coronal slices, slice thickness 280 μm, number of averages 20. Three mice were discarded from the analyses owing to the presence of large susceptibility distortions in the DW images due to the presence of air bubbles following imperfect perfusion procedure. As a result of this, the final number of subjects per group was *n* = 13 and *n* = 10, for *Cntnap2^+/+^* and *Cntnap2^−/−^*, respectively.

#### White-matter fibre tractography

The DW datasets were first corrected for eddy current distortions (FSL/eddy_correct) and skull-stripped (Oguz et al., 2014). The resulting individual brain masks were manually corrected using ITK-SNAP (Yushkevich et al. 2006). Whole brain tractography was performed using MRtrix3 (Tournier et al., 2012) using constrained spherical deconvolution (l_max_ = 8, (Tournier et al., 2007)) and probabilistic tracking (iFOD2) with a FOD amplitude cut-off of 0.2. For each dataset, the whole brain mask was used as a seed, and a total of 100,000 streamlines were generated.

The corpus callosum and cingulum were selected as tracts of interest, given their major cortico-cortical extension and direct involvement in prefrontal-posterior connectivity (Vogt and Paxinos, 2014). The tracts were virtually dissected with waypoint VOIs described in Fig. S3 using TrackVis (http://www.trackvis.org/). Inter-group differences in streamline counts of the tracts were evaluated using a two-tailed Student’s *t*-test (t_21_ > 2.08, p < 0.05). To provide a visual assessment of fibre distribution across groups, voxelwise parametric fibre density maps were generated using DiPy (Garyfallidis et al., 2014), by determining for each voxel the number of subjects in which at least one streamline of the fibre tract of interest passes through the voxel. For visualization purposes, both the dissected tracts and group fibre density maps were transformed to the Allen Mouse Common Coordinate Framework, Version 3 (http://www.brain-map.org/).

#### Rabies virus production and injection

Unpseudotyped recombinant SADΔG-mCherry rabies virus (RV) was produced as described by Osakada and Callaway (2013). Briefly, B7GG packaging cells, which express the rabies envelope G protein, were infected with unpseudotyped SADΔG-mCherry-RV’ obtained by courtesy of Prof. Edward Callaway from the Salk Institute. After five to six days, the virus was collected, filtrated with 0.45 μm filter and concentrated by two rounds of ultracentrifugation. The titer of the SADΔG-mCherry-RV preparation was established by infecting Hek-293T cells (ATCC cat n° CRL-11268) with tenfold serial dilution of viral stock, counting mCherry expressing cells 3 days after infection. The titer was calculated as 2×10^n^ Infective Units (IU)/ml, and the stock was therefore considered suitable for in vivo microinjection. Intracortical rabies virus injections were carried out as previously described (Sforazzini et al., 2014b) in adult (12-16 week-old) male *Cntnap2^−/−^* and control *Cntnap2^+/+^* littermates (*n* = 6 each group). To this purpose, mice were deeply anesthetized with avertin (250 mg/kg) and firmly stabilized on a stereotaxic apparatus (Stoelting Inc.). A micro drill (Cellpoint Scientific Inc.) was used to drill holes through the skull. Injections were performed with a Nanofil syringe mounted on an UltraMicroPump UMP3 with a four channel Micro4 controller (World Precision Instruments), at a speed of 5 nl/s, followed by a 5–10 minutes waiting period, to avoid backflow of viral solution and unspecific labelling. One μl of viral stock solution was injected unilaterally in the left anterior prefrontal cortex using the following coordinates for injections, expressed in mm from bregma: 1.42 from anterior to posterior, 0.3 lateral, −1.6 deep (Paxinos and Franklin, 2011)

#### Quantification of retrogradely labelled cells

RV-labelled cell quantification and histological analyses where carried out by an operator (A.B.) blind to genotype. After 7 days from viral injection, the animals were transcardially perfused with 4% paraformaldehyde (PFA), brains were dissected, post-fixed over night at 4°C and vibratome-cut (Leica Microsystems). RV-infected cells were detected by means of immunohistochemistry performed on every other 100 μm thick coronal section, using rabbit anti-red fluorescent protein (RFP) primary antibody (1:500, AbCam), and goat anti-rabbit HRP secondary antibody (1:500, Jackson ImmunoResearch), followed by 3-3′ diaminobenzidine tetrahydrochloride (DAB, Sigma Aldrich) staining. Imaging was performed with MacroFluo microscope (Leica). Each picture was then superimposed onto the corresponding Paxinos Atlas table (Paxinos and Franklin, 2011), and cell bodies were plotted according to their anatomical localization. The cells were then assigned to their corresponding brain regions, and final region-based cell population counts were expressed as fraction of the total amount of labelled cells.

#### Histological and immunohistochemical analysis of white matter

To histologically assess the presence of microstructural white matter alterations, we examined immunofluorescence-enhanced coronal brain sections covering anterior callosal regions from adult (12 week-old) male *Cntnap2^−/−^* and control *Cntnap2^+/+^* littermates (*n* = *5*, each group) after incubation with rat anti-myelin basic protein (MBP) primary antibody (1:1000, AbCam), followed by donkey anti-rat 594 secondary antibody (1:500, Thermo scientific). We also quantified MBP levels as previously described (Mottershead et al., 2003; Richetto et al., 2016). Briefly, three representative random images in anterior callosal regions characterized by parallel or transversal fiber extension with respect to the image plane (corpus callosum and forceps minor of the corpus callosum, respectively) were acquired on a Nikon A1 confocal system, equipped with 561 laser diode and appropriate filter for Texas Red fluorophore. Z-stack images (1.5 μm thick) were acquired using an oil-immersion 60× plan-apochromat objective at 1024×1024 pixel resolution. Callosal image fields were also qualitatively inspected for the presence of inter-group differences in white matter organization or reduced neuronal packing/density. MBP content was empirically quantified by summing MBP-immunoreactive areas expressed as number of pixels whose values were above the background threshold, calculated as pixel intensity values in areas with no detectable immunostaining, such as cell nuclei or MBP-devoid background.

### Results

#### Reduced local and long-range connectivity in fronto-cortical regions of Cntnap2^−/−^ mice

To obtain an unbiased mapping of genotype-dependent differences in functional connectivity, we implemented recently developed aggregative metrics for local and long-range functional connectivity. This analysis revealed foci of significantly reduced local and long-range connectivity in *Cntnap2^−/−^* mutants with respect to wild-type control subjects (*t*-test, p < 0.05 FWE cluster-corrected, with cluster-defining threshold of *t*_24_ > 2.06, p < 0.05; Fig. 1) encompassing prefrontal (prelimbic and cingulate) and retrosplenial cortices. These same brain regions have been classified both in mice and in humans as “high strength” functional connectivity hubs (Buckner et al., 2009; Cole et al., 2010; Liska et al., 2015), and as such are thought to play a key integrative role in distributed functional networks. Local connectivity reductions appeared to be more widespread than corresponding long-range connectivity deficits (Fig. 1a, c), encompassing involvement of supplementary motor areas surrounding cingulate cortex. The observed local and long-range connectivity reductions were statistically significant also when integrated over a large volume of interest encompassing the whole cingulate cortex (local connectivity: Cg, *t*-test, *t*_24_ = 3.11, p = 0.005, Fig. 1b; long-range connectivity: Cg, *t*-test, *t*_24_ = 2.26, p = 0.03, Fig. 1d).

**Figure 1.**
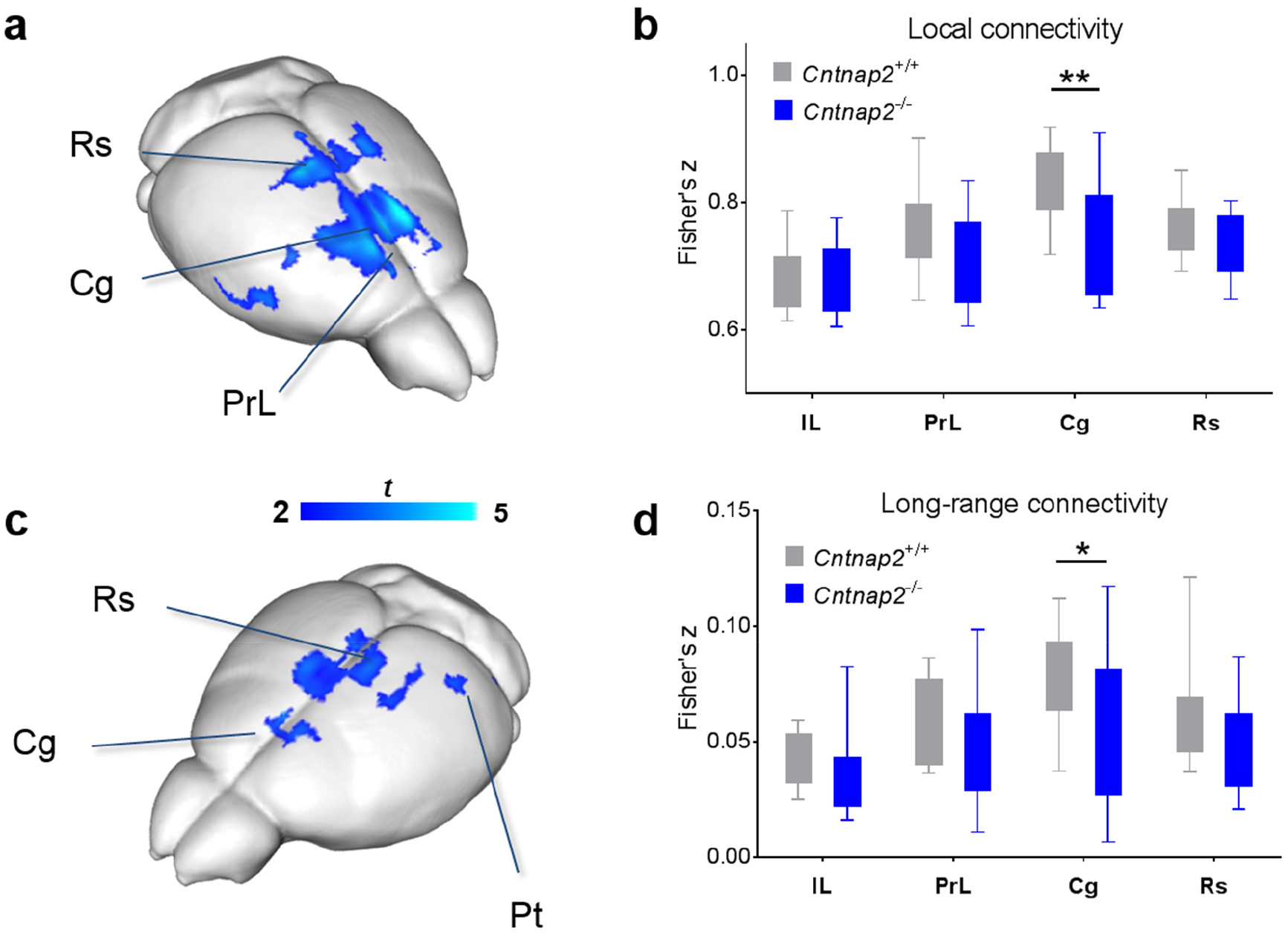
Reduced local and long-range connectivity in Cntnap2^−/−^ mutants. (**a**) Foci of reduced local connectivity in *Cntnap2^-/^* vs. control *Cntnap2^+/+^* littermates (*t*-test; t_24_ > 2.06, p < 0.05; cluster corrected with cluster-level p < 0.05). (**b**) Mean local connectivity in regions of interest (*t*-test; Cg: t_24_ = 3.11, p = 0.005). (**c**) Foci of reduced long-range connectivity in *Cntnap2^-/^* vs. control *Cntnap2^+/+^* littermates (*t*-test; p < 0.05 FWE cluster-corrected, with cluster-defining threshold of t_24_ > 2.06, p < 0.05). (*d*) Mean long-range connectivity in regions of interest (*t*-test; Cg: t_24_ = 2.26, p = 0.03). IL, infra-limbic cortex; PrL, prelimbic cortex, Cg, cingulate cortex; Rs, retrosplenial cortex, * p < 0.05, **p < 0.01.

### Long-range connectivity impairments in Cntnap2^−/−^ mice affect heteromodal cortical regions and the DMN

To identify regional targets of the observed long-range connectivity deficits, we probed rsfMRI networks previously shown to involve prefrontal, cingulate and retrosplenial regions (Sforazzini et al., 2014a; Gozzi and Schwarz, 2015). Seed-based mapping of retrosplenial and anterior cingulate/prelimbic cortex highlighted foci of reciprocal long-range hypoconnectivity along the midline brain axis in *Cntnap2^−/−^* mutants (*t*-test, p < 0.05 FWE cluster-corrected, with cluster-defining threshold of *t*_24_ > 2.06, p < 0.05; Fig. 2a, b). We also probed connectivity of putative lateral components of the rodent DMN such as the posterior parietal and temporal association/auditory cortices, and postero-ventral hippocampus (Gozzi and Schwarz, 2015). Parietal cortical mapping revealed foci of reduced local and long-range (middle cingulate) connectivity in *Cntnap2^−/−^* mice (*t*-test, p < 0.05 FWE cluster-corrected, with cluster-defining threshold of *t*_24_ > 2.06, p < 0.05; Fig. 2c). In the same animals, temporal association areas appeared to be widely hypo-connected to retrosplenial, cingulate and prefrontal regions (*t*-test, p < 0.05 FWE cluster-corrected, with cluster-defining threshold of *t*_24_ > 2.06, p < 0.05; Fig. 2d). We also observed foci of long-range hypo-connectivity between ventral hippocampal and ventral prefrontal (infralimbic) regions (*t*-test, p < 0.05 FWE cluster-corrected, with cluster-defining threshold of *t*_24_ > 2.06, p < 0.05; Fig. 2e). Inter-hemispheric connectivity in subcortical or motor-sensory networks appeared to be overall largely preserved. A reduction in inter-hemispheric connectivity was observed in primary motor areas and visual cortex when quantified in anatomical volumes of interest (Fig. S4), although the effect did not survive false discovery rate correction.

**Figure 2.**
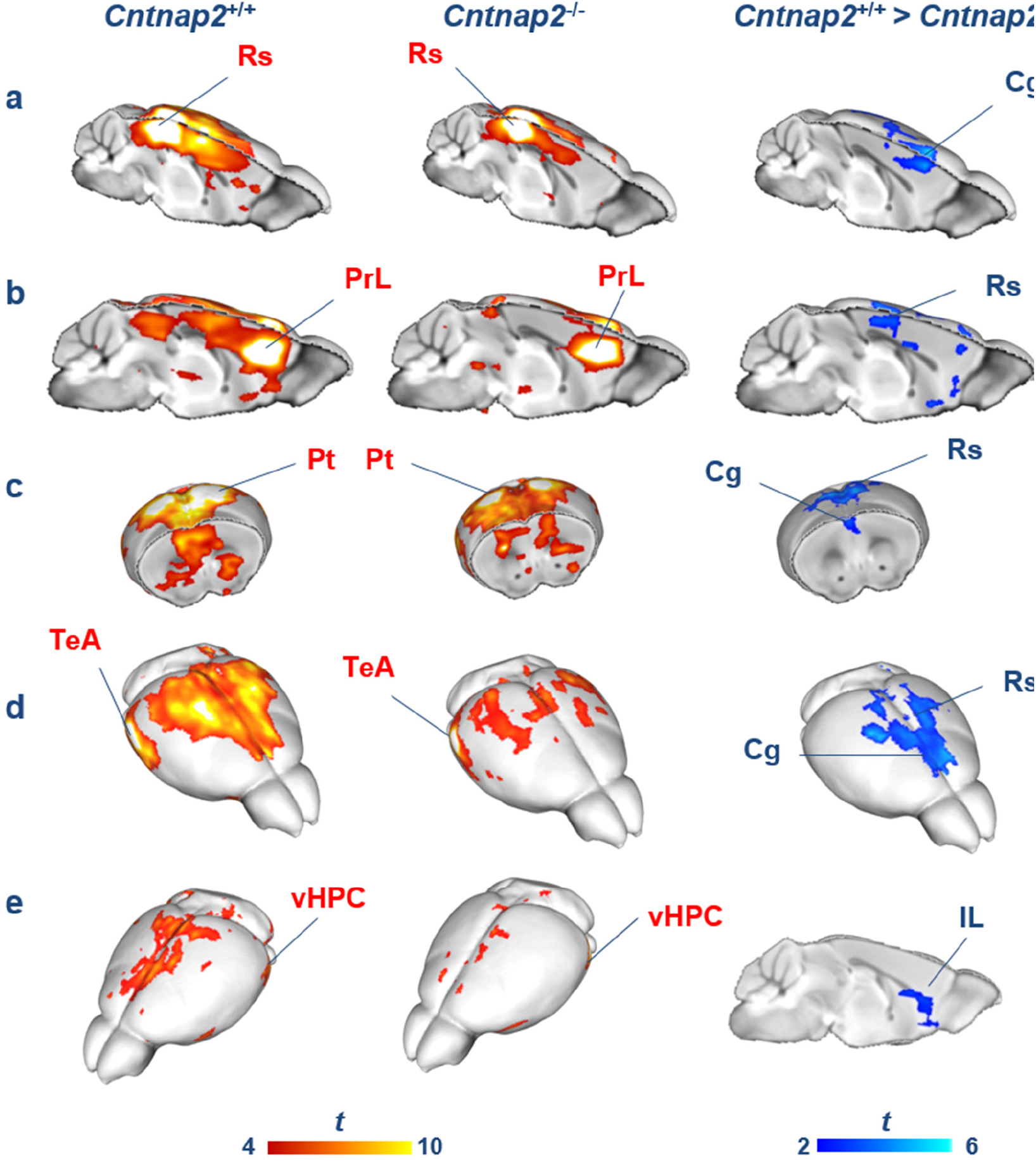
Reduced long-range connectivity in Cntnap2^−/−^ mice. (**a-e**) Seed-correlation mapping highlighted convergent reduced connectivity between long-range cortical and subcortical regions and cingulate-prefrontal areas. Red/yellow shows areas with significant correlation with seed regions indicated in red (one-sample *t*-test, p < 0.05 FWE cluster corrected with cluster-defining threshold of t_12_ > 2.18, p < 0.05). Blue indicates foci of reduced connectivity in *Cntnap2^−/−^* mutants with respect to control mice (*t*-test, p < 0.05 FWE cluster corrected with cluster-defining threshold of *t*_24_ > 2.06, p < 0.05). Rs, retrosplenial cortex; IL, infra-limbic cortex; PrL, prelimbic cortex; Cg, cingulate cortex; Rs, retrosplenial cortex; vHPC, ventral hippocampus; Au/TeA, auditory/temporal association cortices; Pt, parietal cortex.

Importantly, no genotype-dependent differences in anaesthesia sensitivity were detected as seen with mean arterial blood pressure mapping (*t*-test, *t*_24_ = 0.17, p = 0.87; Fig. S5a) and amplitude of cortical BOLD signal fluctuations (*t*-test, *t*_24_ = 0.72, p = 0.48; Fig. S5b), two independent readouts previously shown to be linearly correlated with anaesthesia depth (Steffey et al., 2003; Liu et al., 2011). Together with the observation of region-dependent alterations, as opposed to the global reduction described with increased anaesthesia dosing (Nasrallah et al., 2014), these findings strongly argue against a confounding contribution of anaesthesia to the observed hypo-connectivity.

### Hypoconnectivity in the mouse DMN is associated with impaired social behaviour

Recent human imaging studies in socially-impaired patients have revealed a putative association between long-range DMN hypo-connectivity and social competence (Schreiner et al., 2014). Based on these findings, we hypothesized that reduced long-range DMN connectivity in *Cntnap2^−/−^* mice could be associated with impaired social behaviour. To test this hypothesis, we first corroborated DMN hypoconnectivity by quantifying functional connectivity along the dorsal midline axis of this network (anterior/middle cingulate cortex and retrosplenial cortex) using multiple seed-to-VOI measurements (Fig. 3). A clear dysconnection between posterior (retrosplenial) and middle/anterior portions of the DMN (cingulate, prelimbic cortex) was apparent (retrosplenial to cingulate cortex: two-way repeated-measures ANOVA, genotype effect, F_1,24_ = 5.76, p = 0.02, Fig. 3a; retrosplenial-cingulate to prelimbic: two-way repeated-measures ANOVA, genotype effect, F_1,24_ = 6.82, p = 0.02, Fig. 3b).

**Figure 3.**
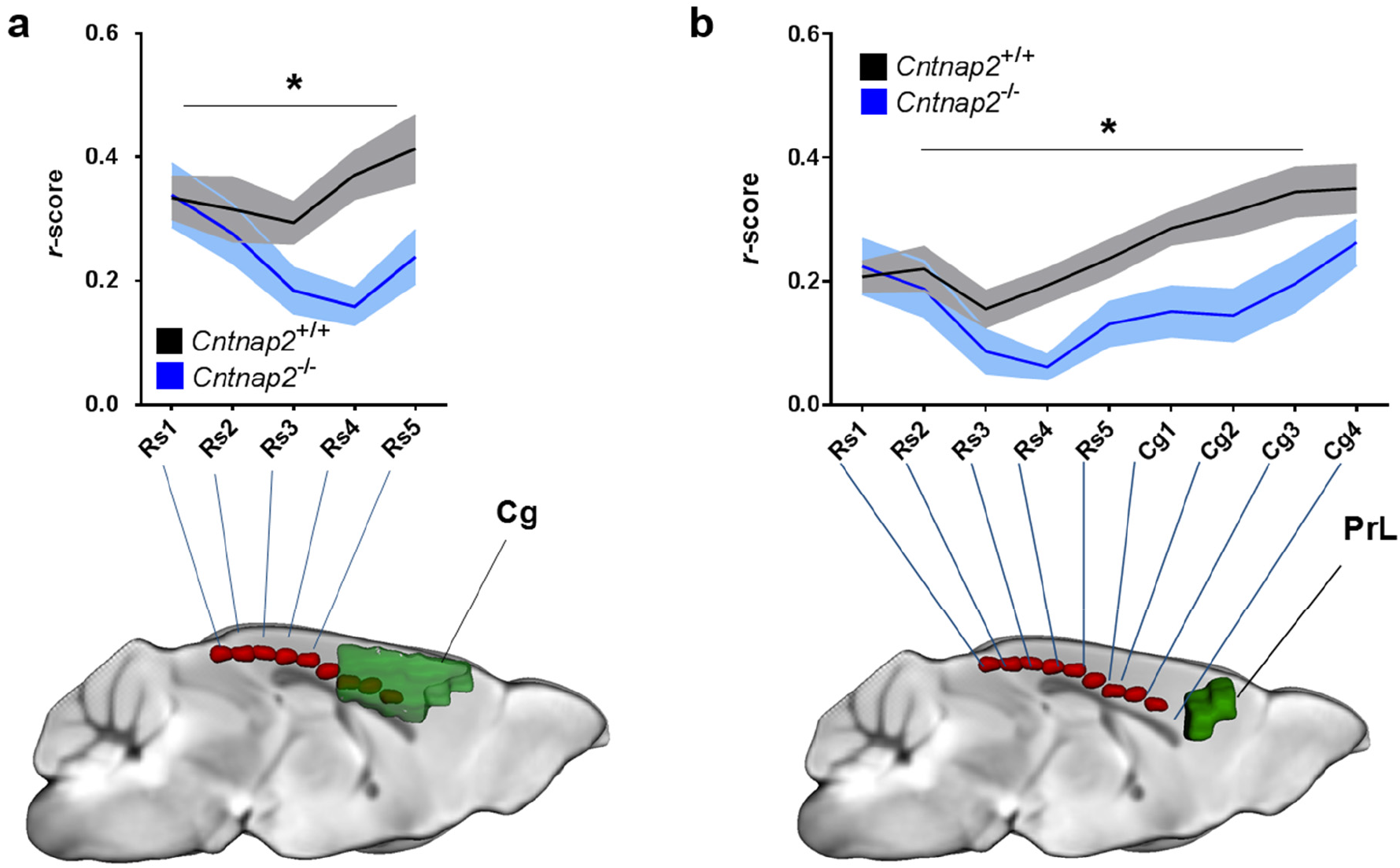
Fronto-posterior hypoconnectivity in Cntnap2^/-^ mice. (**a**) Connectivity profile between a series of retrosplenial seeds (Rs, red) and the cingulate cortex (Cg, green) (two-way repeated-measures ANOVA, genotype effect, F_1,24_ = 5.76, p = 0.02). (**b**) Connectivity profile between a series of retrosplenial/cingulate seeds (Rs, Cg, red) and the prelimbic cortex (PrL, green) (two-way repeated-measures ANOVA, genotype effect, F**1,24** = 6.82, p = 0.02). * p < 0.05

We then measured social behaviour in adult *Cntnap2^−/−^* and *Cntnap2^+/+^* control mice in a male-female interaction test, and correlated the measured social scores with DMN hypoconnectivity measures. Consistent with previous reports (Penagarikano et al., 2011), behavioural testing revealed significantly impaired social interest (total sniffing, duration: *t*-test, *t*_24_ = 2.29, p = 0.03, Fig. 4a; social investigation, duration: *t*-test, *t*_24_ = 2.43, p = 0.02, Fig. 4c) and increased non-social behaviour (wall-rearing, frequency: *t*-test, *t*_24_ = 3.09, p = 0.01; Fig. S6a) in *Cntnap2^−/−^* mutants compared to *Cntnap2^+/+^* control littermates. Hypo-connectivity in key DMN components (retrosplenial-cingulate cortex) was significantly associated with reduced social behaviour (total sniffing, duration: r = 0.42, p = 0.03, n = 26, *R*^2^ = 0.17, Fig. 4b; social investigation, duration: r = 0.40, p = 0.04, n = 26, *R*^2^ = 0.16, Fig. 4d) and increased non-social behaviour (wall rearing, frequency: r = −0.45, p = 0.02, n = 26, *R*^2^ = 0.21; Fig. S6b). These findings highlight a correlation between fronto-posterior connectivity and social behaviour, suggesting that impaired functional couplings produced by mutations in *Cntnap2* could reverberate to affect complex behavioural traits such as sociability and social exploration.

**Figure 4.**
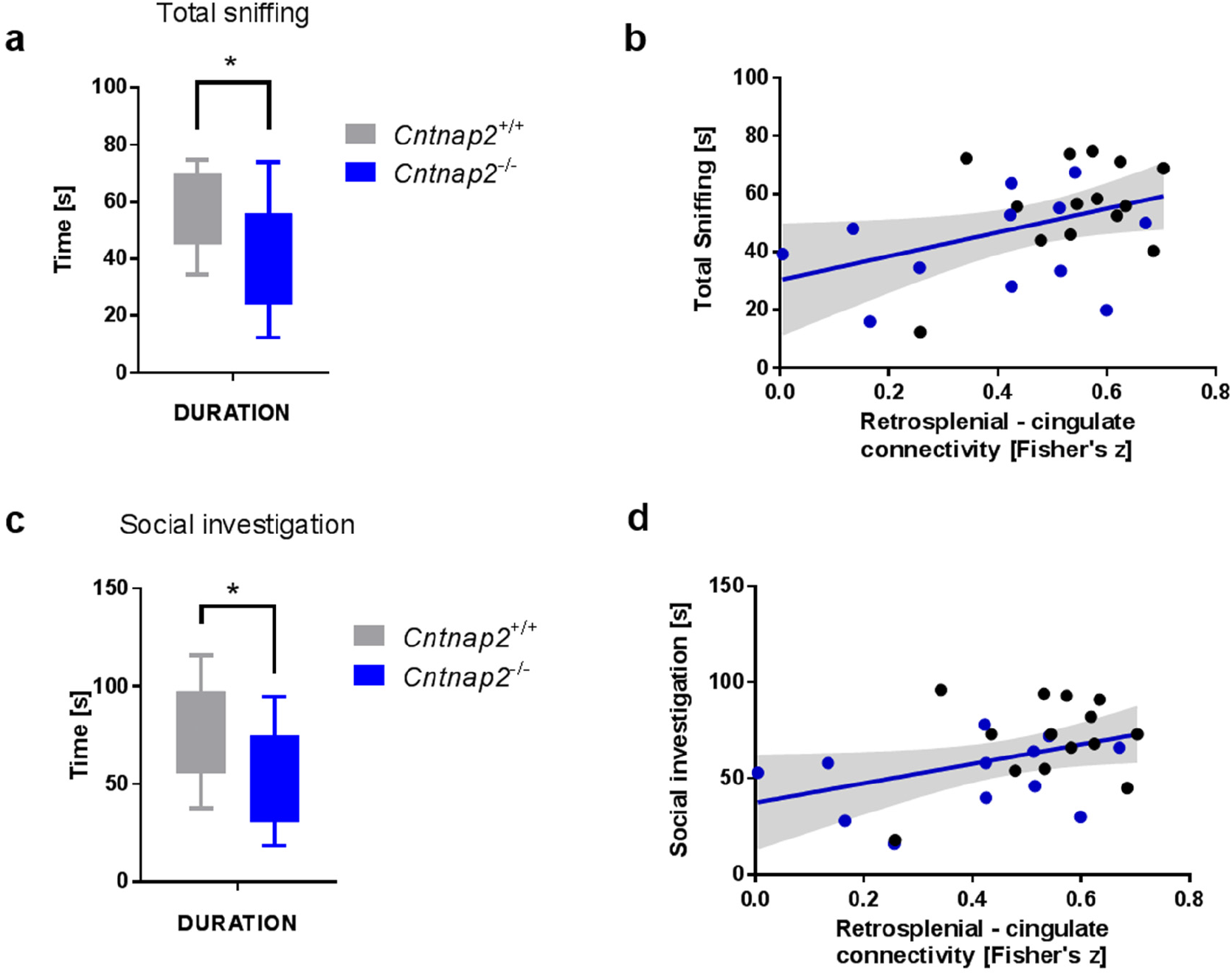
Fronto-posterior connectivity is correlated with social behaviour. (**a**) Social behaviour as measured by total sniffing duration (*t*-test, t_24_ = 2.29, p = 0.03). (**b**) Association between retrosplenial-cingulate connectivity (VOI to VOI) and total sniffing duration (r = 0.42, p = 0.03, n = 26, *R*^2^ = 0.17). (**c**) Social behaviour as measured by the duration of social investigation (*t*-test, t_24_ = 2.43, p = 0.02). (d) Association between retrosplenial-cingulate connectivity (VOI-to-VOI) and the duration of social investigation (r = 0.40, p = 0.04, n = 26, *R*^2^ = 0.16). * p < 0.05.

### Macroscale cortico-cortical white matter connectivity is preserved in Cntnap2^−/−^ mice

To probe a role of *macroscale* anatomical connectivity alterations on the observed functional decoupling in *Cntnap2^−/−^*, we performed tractography analysis of the corpus callosum and cingulum, two major white matter tracts characterised by extensive cortico-cortical antero-posterior extension (Fig. 5a). These white matter tracts appeared to be largely typical in mutant and control mice as seen with group-level fibre density maps (Fig. 5b); in keeping with this, we did not observe statistically significant differences in the number of streamlines between *Cntnap2^−/−^* mutants and controls (cingulum: *t*-test, t_21_ = 1.25, p = 0.23; corpus callosum: *t*-test, t_21_ = 1.21, p = 0.24; Fig S7). These results argue against a contribution of gross macroscale white matter alterations to the observed functional connectivity impairments.

### Reduced prefrontal-projecting neuronal clusters in cingulate cortex of Cntnap2^/-^ mice

Although macroscale cortico-cortical connectivity appeared to be normal in Cntnap2^−/−^ mutants, the possibility exists that finer-scale miswiring, undetectable by tractography, may contribute to the mapped functional connectivity alterations. To probe this hypothesis, we carried out monosynaptic retrograde tracing of the left prefrontal cortex (prelimbic/anterior cingulate cortex area 1, corresponding to Brodmann area 24Ab, Vogt and Paxinos, 2014) and quantified the number of retrogradely labelled cells in representative volumes of interest in mutant and control littermate mice (Fig. 6a). The anatomical distribution of retrogradely labelled neurons in both genotypes was in keeping with previously published rodent studies (Hoover and Vertes, 2007) and encompassed several key anatomical substrates considered to be part of the rodent DMN (Gozzi and Schwarz, 2015). Notably, regional quantification of the relative fraction of labelled cells revealed reduced frequency of prefrontal-projecting neurons in the cingulate cortex of *Cntnap2^−/−^* mutants (Cg: *t*-test, t_10_ = 3.90, p = 0.003, FDR-corrected p = 0.04; Fig. 6b, c). Importantly, no genotype-dependent significant difference in the number of prefrontal projecting neurons was observed in any of the other cortical or subcortical regions examined (Fig. 6c).

**Figure 5.**
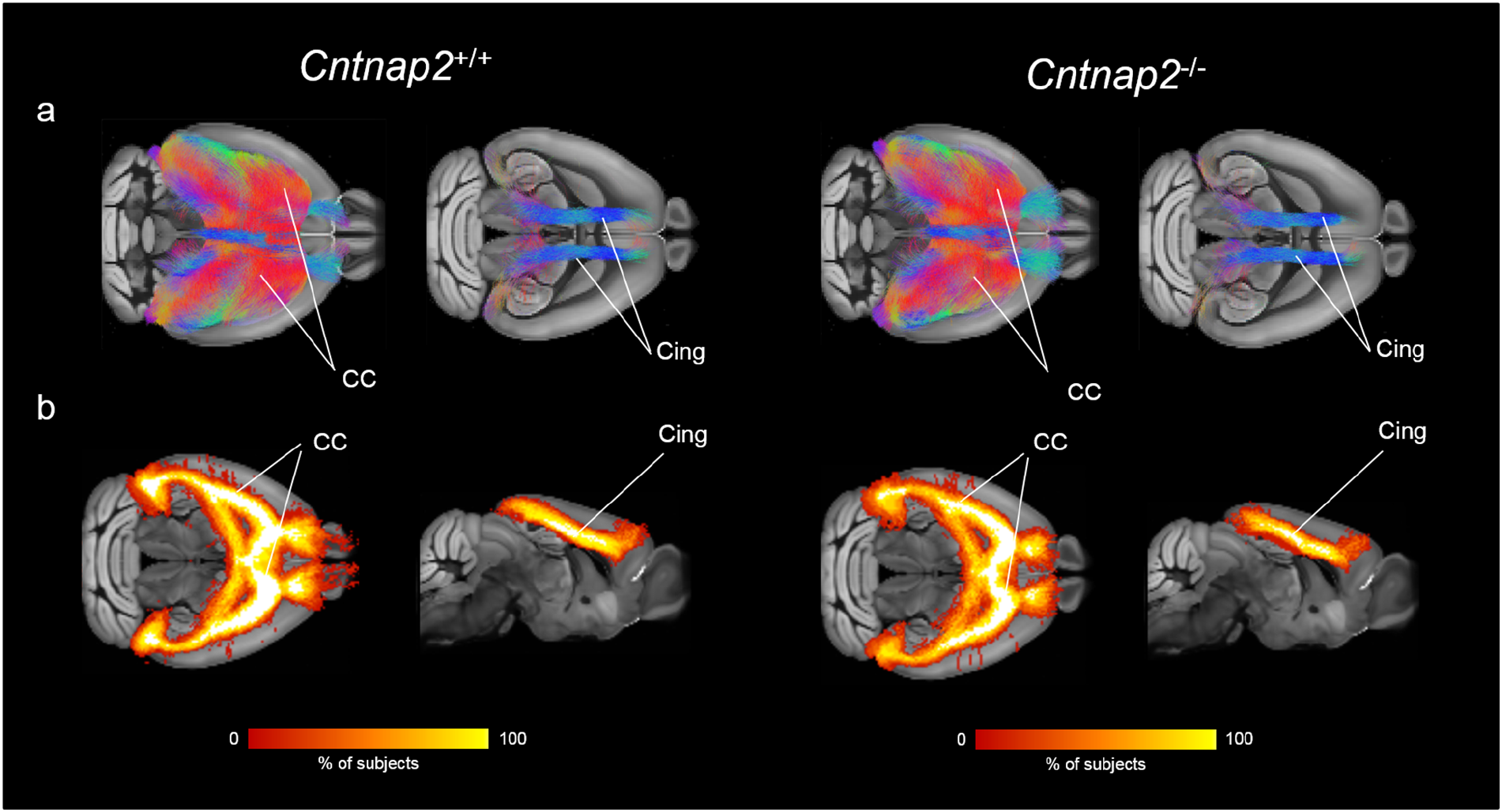
Preserved cortico-cortical white matter organization in Cntnap2^−/−^ mutants. (**a**) Corpus callosum and cingulum tracts virtually dissected in two representative subjects (*Cntnap2^+/+^* left, *Cntnap2^−/−^* right), (**b**) Fractional group fibre density maps for corpus callosum and cingulum tracts (*Cntnap2^+/+^* left, *Cntnap2^−/−^*, right).

**Figure 6.**
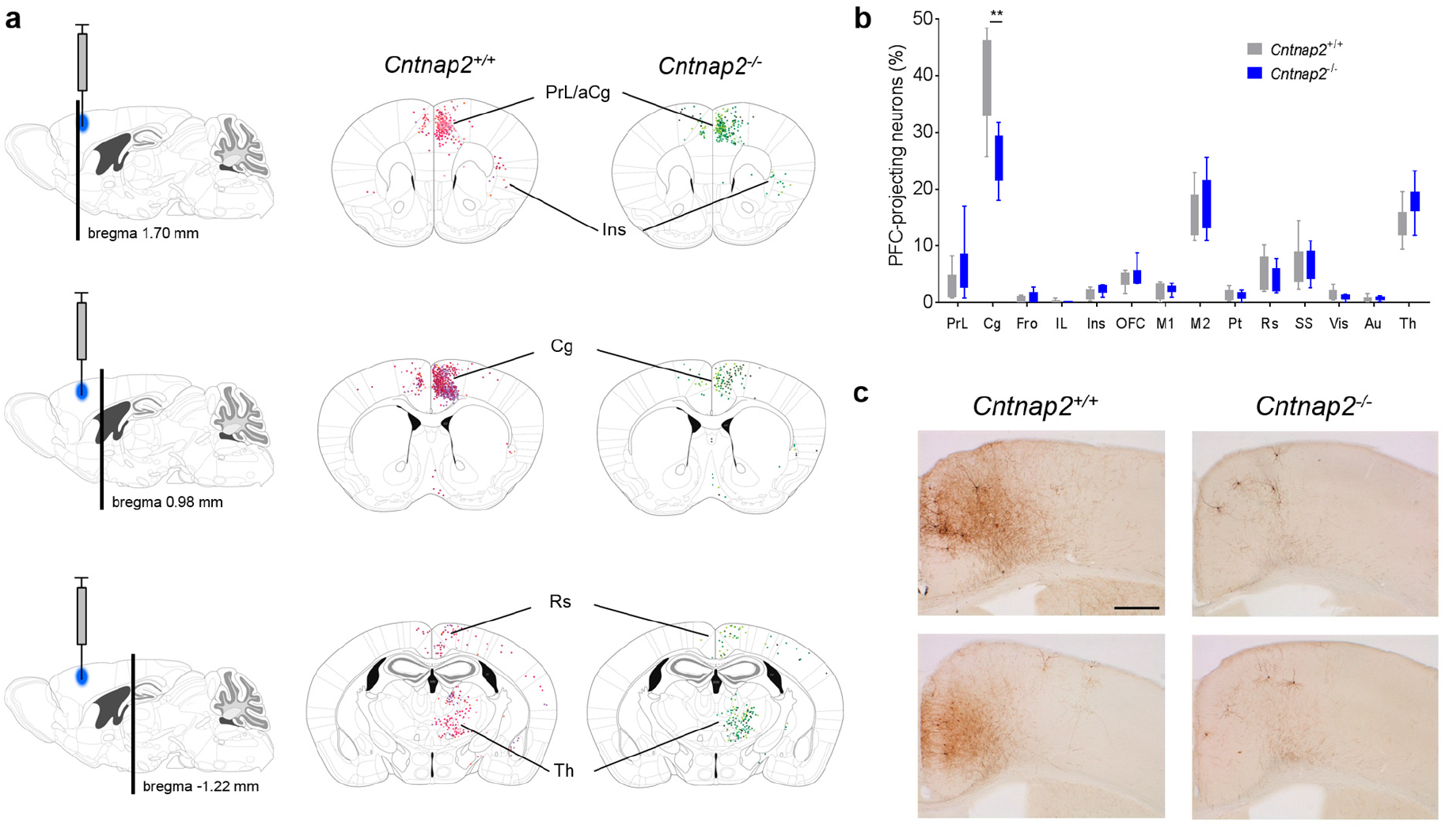
Reduced frequency of cingulate-prefrontal projecting neurons in Cntnap2^−/−^ mice. (**a**) Locations of retrogradely labelled cells superimposed on the corresponding Paxinos Atlas coronal tables. Injection location is indicated in blue on the sagittal tables. (**b**) Regional quantification of the relative regional number (frequency) of retrogradely labelled cells (*t*-test; Cg: t_10_ = 3.90, p = 0.003, FDR-corrected p = 0.04). (**c**) Enlarged view of the distribution of retrogradely labelled cells in a coronal section of the cingulate region (bregma 0.98 mm) in two representative *Cntnap2^+/+^* and two *Cntnap2^−/−^* subjects. The scale bar indicates 250 μm. * p < 0.05, ** p < 0.01.

### Preserved microscale white matter organization in Cntnap2^/-^ mice

We next examined the presence of *microscale* white matter structural abnormalities in control and *Cntnap2^−/−^* mutants via histological examinations and myelin binding protein (MBP) quantification. In keeping with previous investigations ((Poliak et al., 2003; Penagarikano et al., 2011), we did not observe gross microscale white matter disorganization or morphological changes in mice lacking *Cntnap2* with respect to control littermates (Fig. S8a). Similarly, MBP quantification in frontal callosal white matter tracts did not reveal any significant between-group difference (corpus callosum: *t*-test, t_8_ = 0.84, p = 0.42; forceps minor of the corpus callosum: *t*-test, t_8_ = 1.06, p = 0.32; Fig. S8b).

## Discussion

Here we show that homozygous mice lacking *Cntnap2*, a neurexin superfamily member associated with autism, exhibit reduced long-range and local functional connectivity in prefrontal cortical regions and midline functional hubs of the mouse brain, an effect that may involve defective cingulate-prefrontal mesoscale wiring. We also show that reduced fronto-posterior connectivity is associated with impaired social behaviour, revealing a possible link between long-range functional connectivity alterations and mouse behavioural traits recapitulating ASD symptoms. Collectively, these findings suggest that loss-of-function mutations in *Cntnap2* may predispose to neurodevelopmental disorders and autism through dysregulation of macroscale functional network couplings.

Our use of an imaging readout widely employed in human connectivity mapping provides us with the opportunity to cross-compare connectivity findings across species. In this respect, the observation of long-range fronto-posterior hypo-connectivity in *Cntnap2^−/−^* mice is especially noteworthy, because it is in excellent agreement with the results of a recent human imaging study where an association between common genetic variants in *CNTNAP2* and similar long-range frontal hypoconnectivity was described (Scott-Van Zeeland et al., 2010). Our results expand these findings, by revealing a causal contribution of *Cntnap2* loss-of-function mutations to long-range fronto-cortical connectivity impairments. These correspondences also serve as an important proof-of-concept demonstration that ASD-related genetic mutations can lead to comparable macroscale connectivity deficits in humans and lower mammal species like the laboratory mouse. The presence of long-range hypoconnectivity in *Cntnap2^−/−^* mice also adds to the remarkable construct and face validity of this mouse model as an experimental tool for mechanistic and therapeutic investigation of syndromic forms of ASD (Penagarikano et al., 2011). Specifically, *Cntnap2^−/−^* mice closely recapitulate major neuropathological features observed in cortical dysplasia-focal epilepsy (CDFE) syndrome, a rare neuronal migration disorder associated with a recessive (suggesting loss of function) mutation in *CNTNAP2*, and, in nearly two thirds of patients, with autism (Strauss et al., 2006). These include behavioural deficits in the three core domains of ASD (reduced vocal communication, repetitive and restricted behaviours, and abnormal social interactions), hyperactivity and epileptic seizures (both features described in CDFE patients), and reduced GABAergic interneurons, resulting in asynchronous cortical activity as measured with in vivo two-photon calcium imaging (Penagarikano et al., 2011).

The observation of defective mesoscale axonal wiring in the cingulate cortex corroborates the presence of selective prefrontal dysregulation in *Cntnap2^−/−^* mutants, and serves as a possible neuroanatomical correlate for some of the prefrontal functional connectivity impairments mapped with rsfMRI. Regional differences in GABAergic interneuron density, and developmental processes related to circuit and network refinements (Zhan et al., 2014; Riccomagno and Kolodkin, 2015) are, however, likely to play a role in the observed functional desynchronization as well, given the established contribution of GABAergic oscillatory rhythms in mediating large-scale functional synchrony (Gonzalez-Burgos and Lewis, 2008) and the recent evidence of altered spine density and increased spine eliminations in *Cntnap2^−/−^* mice (Gdalyahu et al., 2015). The relative contribution of anatomical versus neurophysiological mechanisms in determining the observed desynchronization remains however undetermined, and interventional studies entailing the regional manipulation of excitatory/inhibitory ratio or inactivation of projection-specific pathways may be required to disambiguate this issue.

Recent human studies described possible microstructural white matter alterations in carriers of *CNTNAP2* mutations as assessed with water diffusion anisotropy. Specifically, gender-dependent reductions in fractional anisotropy (FA) in the inferior fronto-occipital fasciculus or anterior thalamic radiation have been described by Tan and colleagues (2010). Similarly, Clemm von Hohenberg (2013) described an interaction between a single genotype (rs2710126) and FA, in which homozygotes for the risk allele showed reduced FA values in uncinate fasciculus.

These preliminary results suggest the presence of possible white matter microstructural alterations as a result of *CNTNAP2* gene mutations. However, anisotropic water diffusion reflects multiples biophysical contributions that prevent an unequivocal microstructural interpretation of these findings. For example, reduced FA could be the result of reduced neuronal packing, myelinisation, axonal diameter, neuronal integrity and maturation, as well as regional differences in gray matter fraction (Beaulieu, 2009). To investigate potential microstructural white matter disruption at a more detailed level than permitted by diffusion MRI, we carried out a histological assessment of white matter fibres using MBP immunofluorescence. We did not observe any gross white matter microstructural abnormality in *Cntnap2^−/−^* mutants in terms of fibre orientation, packing or organization. Moreover, MBP quantification did not reveal any significant genotype-dependent differences. Together with previous electron microscopy investigations, where normal myelin thickness was reported in *Cntnap2^−/−^* mutants (Poliak et al., 2003), these findings argue against the presence of gross microscale white matter alterations in these mutants. While this finding appears to be in contrast with human investigations of *CNTNAP2* polymorphisms and suggestive of possible species-specific divergence, additional research is required to more thoroughly investigate the presence of white matter microstructural aberrancies in *Cntnap2^−/−^* mutants. It should also be noted that *CNTNAP2* polymorphisms studies in humans are typically correlative and involve small patient samples, which make them more prone to confounding factors related to heterogeneity in clinical samples, and individual adaptive differences in microstructural parameters (Scholz et al., 2009).

The observation of hypoconnectivity in prefrontal hub regions of the DMN (Liska et al., 2015) is suggestive of a deficient “maturation” of this functional network (Supekar et al., 2010), and is in keeping with the hypothesis of a key role of this region as a mediator of deficits in global perception and its cognitive representations in ASD patients (Martinez-Sanchis, 2014). The notion that “underconnectivity” may preferentially affect complex cognitive and social functions and their high order cortical substrates rather than low-level sensory and perceptual tasks has recently found some theoretical support (Kana et al., 2011). Within this framework, heteromodal integrative hubs like the anterior cingulate and prefrontal cortex, as well as retrosplenial regions would serve as major points of vulnerability for the stability of distributed functional network couplings. rsfMRI mapping in additional mouse lines harbouring ASD-related genetic mutations will be instrumental in assessing whether the observed alterations represent a generalizable endophenotype that may converge across mutations and genetic etiologies, or are the specific consequence of *Cntnap2* mutations. It is, however, interesting to note that so far hypoconnectivity appears to be predominant in mouse imaging studies of ASD: reduced connectivity in several brain regions including the prefrontal cortex and the DMN has been observed in the BTBR model of idiopathic autism (Sforazzini et al., 2014b), and in mice characterised by reduced synaptic pruning (Zhan et al., 2014), a pathological trait associated with autism (Tang et al., 2014). Reduced connectivity between motor sensory regions and a general reduction in primary visual cortex connectivity were also recently described in a mouse model of fragile × syndrome (Haberl et al., 2015). Although preliminary, these initial mouse findings are consistent with and somehow support the “under-connectivity theory” of autism, according to which reduced functional connectivity, at least in the adult brain (Uddin et al., 2013), may emerge as a dominant feature of ASD in the face of heterogeneous etiopathological pathways (Di (Martino et al., 2013; Uddin et al., 2013).

In contrast with our imaging results, human rsfMRI mapping in *CNTNAP2* common variant carriers revealed increased, instead of decreased, local connectivity in lateral prefrontal regions (Scott-Van Zeeland et al., 2010). The reason behind this discrepancy is not clear, although several important experimental factors, including methodological, species- and/or age-related differences may contribute to this inconsistency. For example, local connectivity was found to be increased in human lateral prefrontal areas, a region that does not have a clear cyto-architectural correlate in rodents (Vogt and Paxinos, 2014). Moreover, our study was performed in adult male subjects, while human mapping was carried out in pre-pubertal subjects (mean age 12 years old), a discrepancy that could account for the differences in local connectivity alterations. Indeed, a dramatic reorganization of large-scale functional brain networks occurs during childhood and late adolescence in humans, involving developmental shifts from short-range to long-range connectivity, notably within fronto-insular and cortico-subcortical networks (reviewed by Ernst et al., 2015). Moreover, an age-related dichotomy has been suggested in ASD-related connectivity aberrancies, with generally reduced intrinsic functional connectivity in adolescents and adults with autism compared with age-matched controls, and increased functional connectivity in younger children with the disorder (Uddin et al., 2013). A similar age-related shift could therefore possibly explain the discrepant direction of local connectivity observed in our study with respect to the finding reported by Van Zeeland et al., (2010) in human pre-adolescent subjects. Longitudinal investigations of connectivity in rodent genetic models of autism are highly warranted to enable empirical testing of the developmental trajectory of ASD-related connectivity aberrancies across development and network maturation (Liska and Gozzi, 2016). Finally, differences in the nature of the investigated mutations should also not be neglected, as the functional consequences of the genetic variants imaged by Scott-Van Zeeland et al. (2010) are unclear, and the possibility that not all the imaged genetic variants are loss-of-function cannot be ruled out.

We also note here that our study specifically addressed male mice only, owing to the greater ASD incidence in this gender (Lai et al., 2015). While this choice has the advantage of reducing within-group variation and subsequent increase in statistical power of our measurements, this should be considered a limitation of our study, as it does not permit to assess whether our findings can be generalized to female carriers of *Cntnap2* loss-of-function mutations.

In conclusion, we document that the absence of *Cntnap2* leads to functional connectivity reductions and defective mesoscale wiring in prefrontal functional hubs of the mouse brain, an effect associated with impaired social behaviour. These findings suggest that loss-of-function mutations in *Cntnap2* may predispose to neurodevelopmental disorders and autism through selective dysregulation of connectivity in integrative prefrontal areas, and provide a translational model for investigating connectional perturbations in syndromic ASD forms.

## Acknowledgments

The study was funded by grants from the Simons Foundation (SFARI 314688 and 400101, A.G.).

### Conflict of interests

The authors declare that they have no conflict of interest.

## 5 Supplementary figures

**Figure S1.**
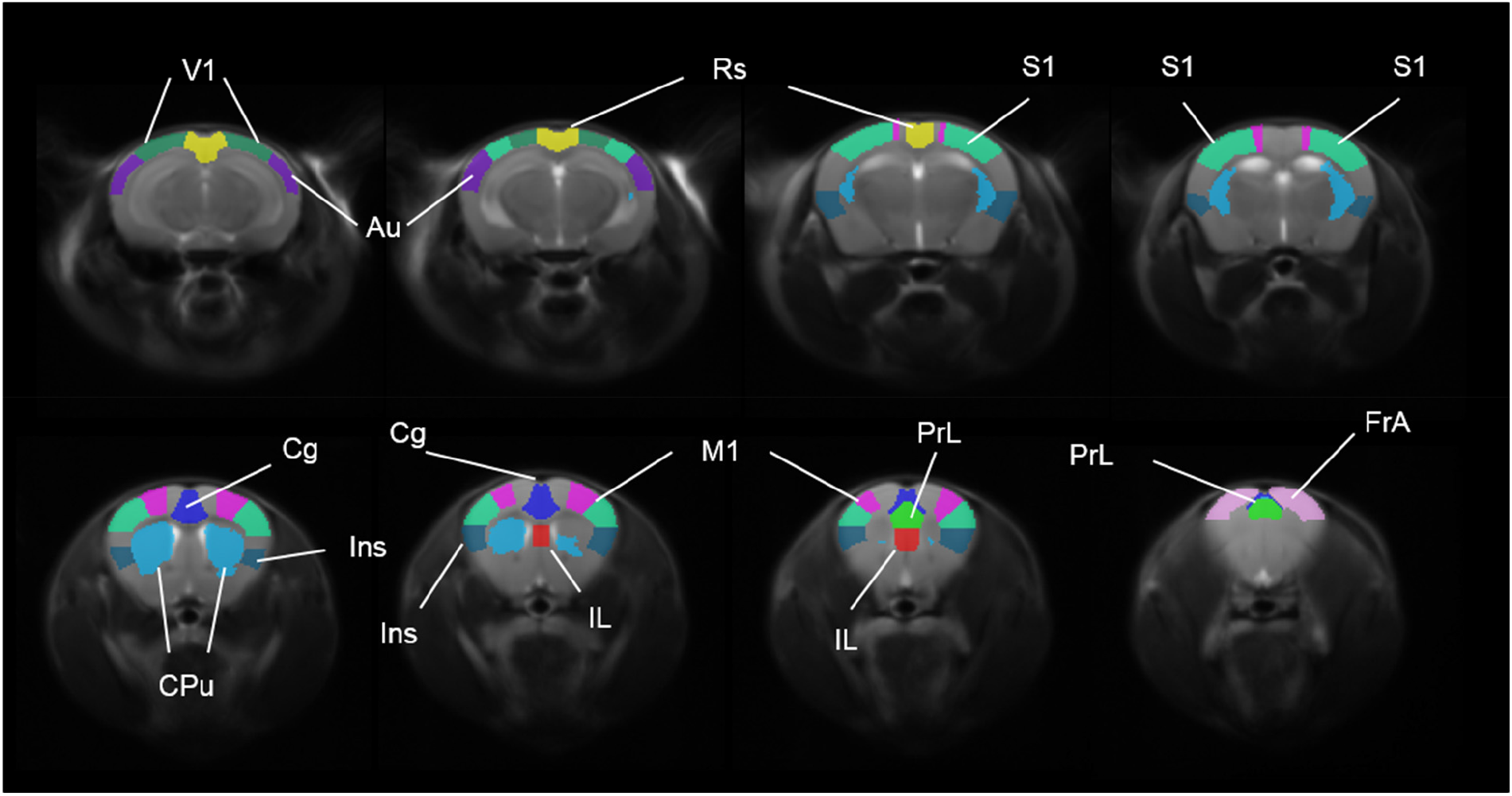
Volumes of interest used in functional connectivity mappings. V1, primary visual cortex; Au, auditory cortex; Rs, retrosplenial cortex; S1, primary somatosensory cortex; Cg, cingulate cortex; CPu, caudate-putamen; Ins, insular cortex; IL, infra-limbic cortex; M1, primary motor cortex; PrL, prelimbic cortex; FrA, frontal association cortex.

**Figure S2.**
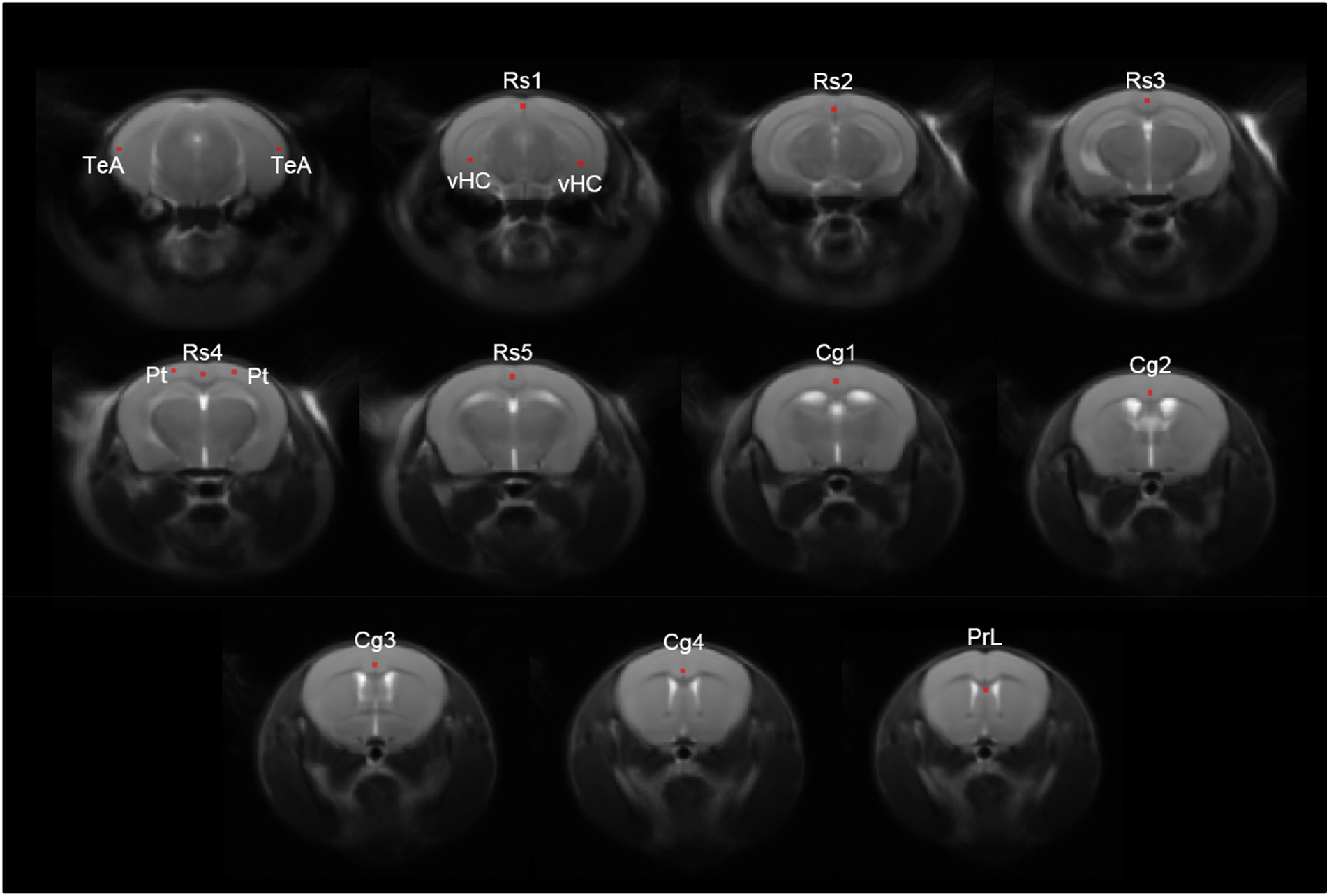
Location of seeds used in mapping anteroposterior DMN connectivity. TeA, temporal association cortex (bilateral); Rs, retrosplenial cortex; Cg, cingulate cortex; Pt, posterior parietal association cortex (bilateral); PrL, prelimbic cortex; vHC: ventral hippocampus.

**Figure S3.**
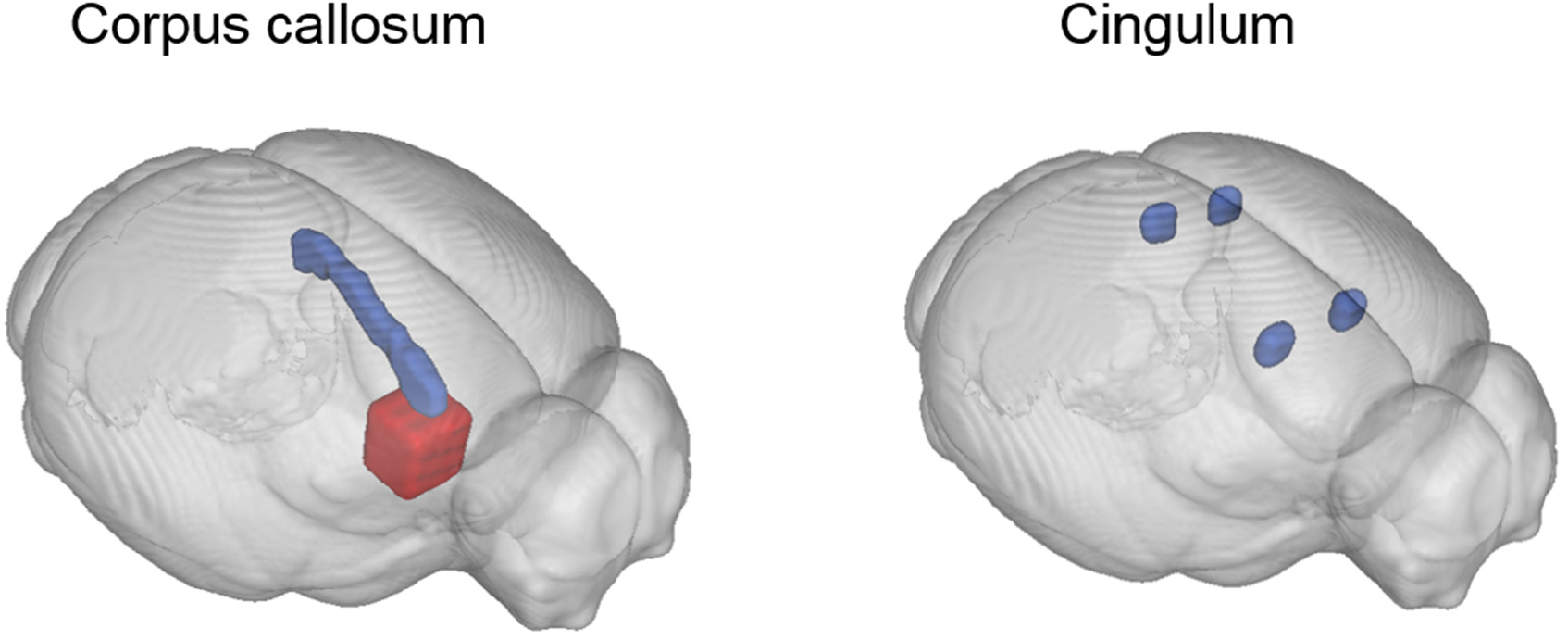
Location of waypoint ROIs used for virtual dissection of corpus callosum and cingulum tracts from whole-brain white matter tractography. Inclusion ROIs are indicated in blue, exclusion ROIs are indicated in red.

**Figure S4.**
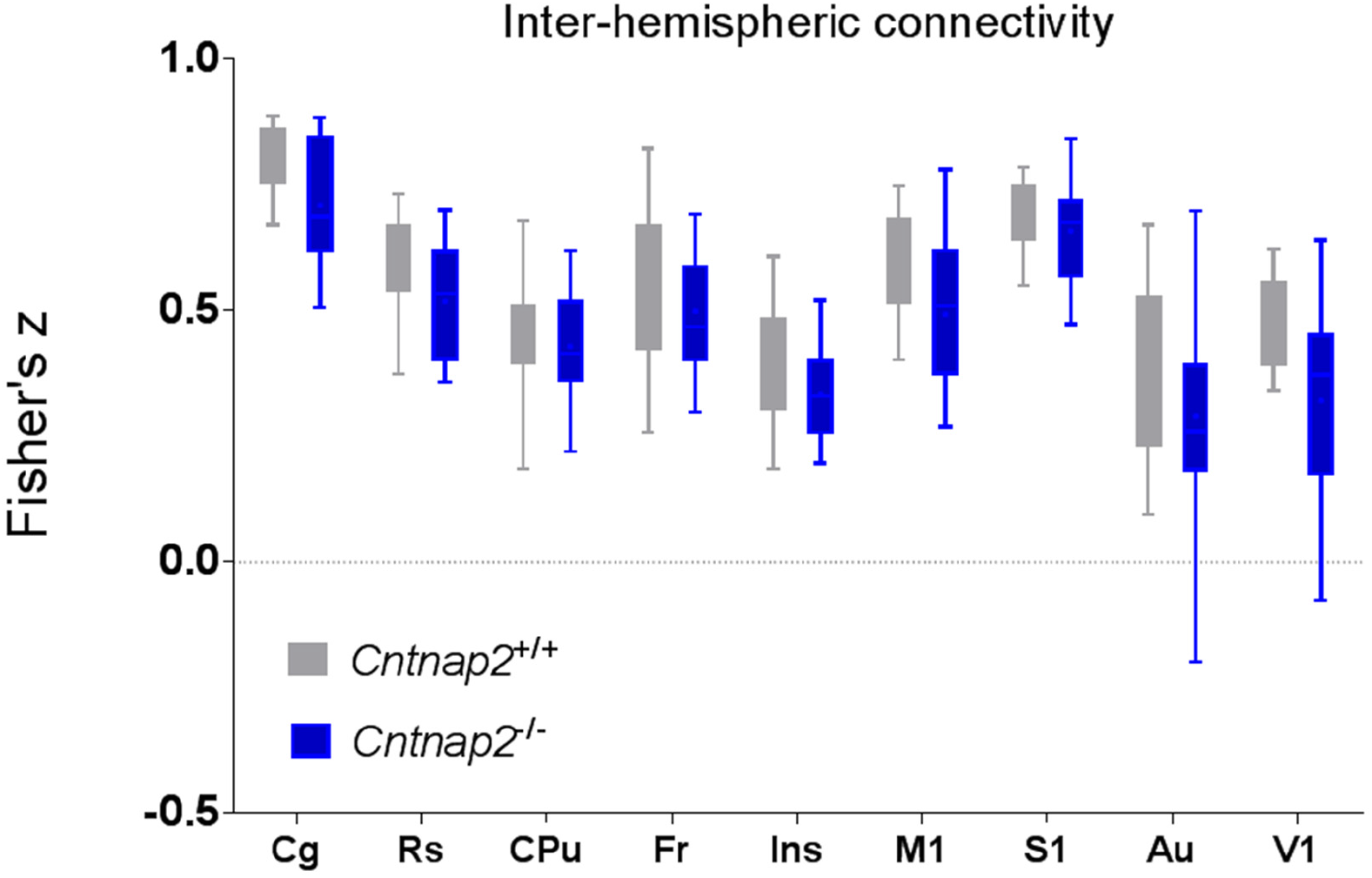
Largely preserved inter-hemispheric connectivity in Cntnap2^−/−^ mutants and control mice. Correlation coefficients were calculated between time courses extracted from VOIs depicted in Fig. S1 and the resulting *r*-scores were transformed to *z*-scores using Fisher’s *r*-to-*z* transform. None of these comparisons survived a false discovery rate correction at q = 0.05.

**Figure S5.**
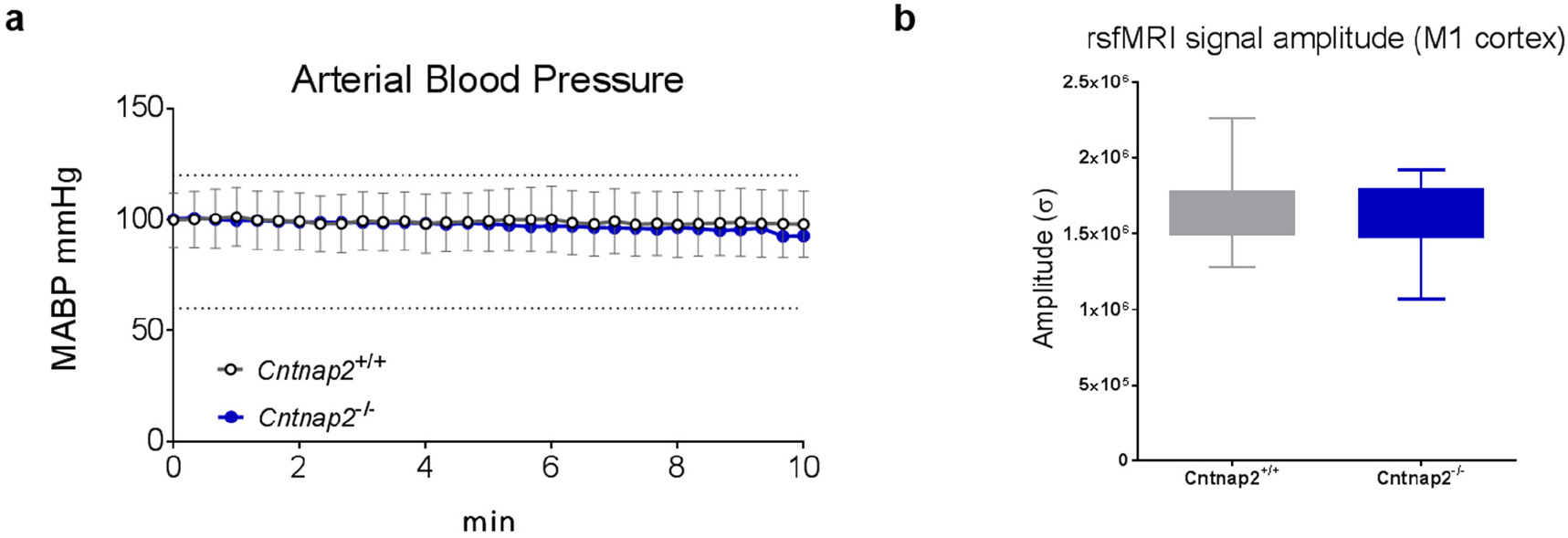
No genotype-dependent differences in anaesthesia sensitivity were detected as seen with mean arterial blood pressure mapping (*t*-test, t_24_ = 0.17, p = 0.87; **a**) and amplitude of cortical BOLD signal fluctuations in primary motor cortex (*t*-test, t_24_ = 0.72, p = 0.48; **b**). M1, primary motor cortex.

**Figure S6.**
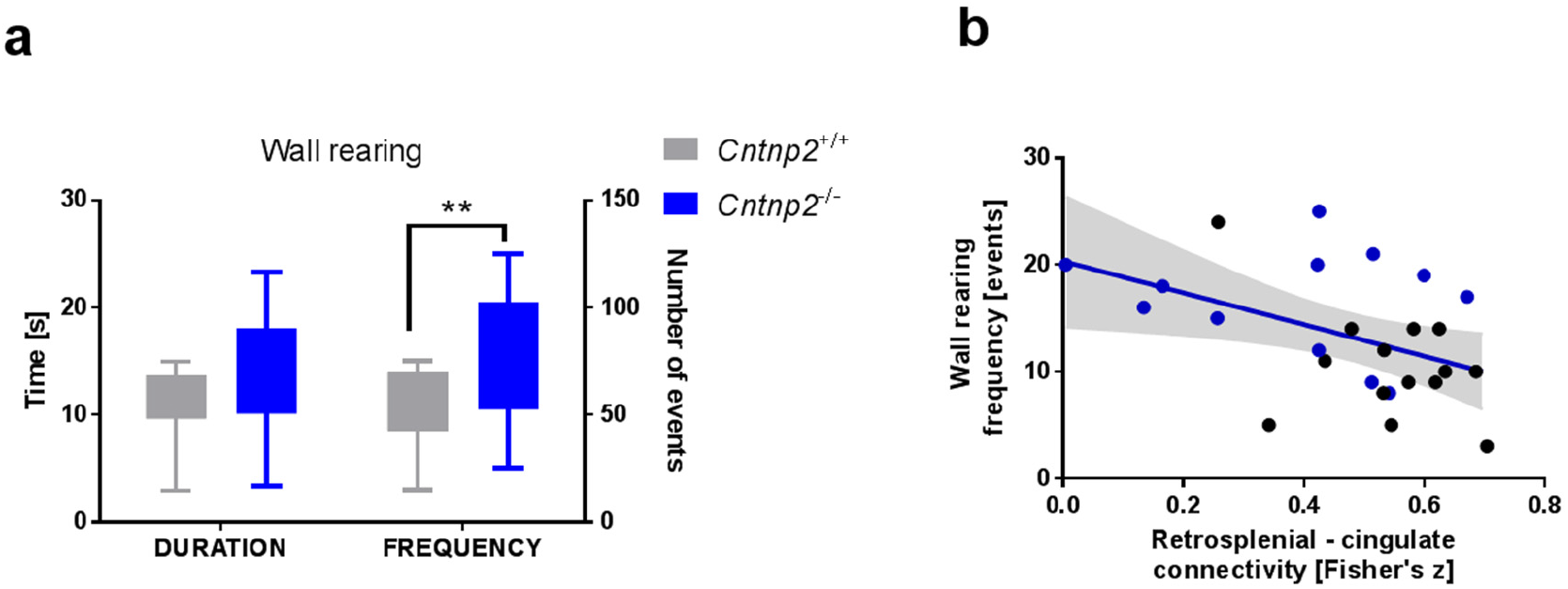
Increased non-social behaviour in Cntnap2^/-^ mutants compared to control littermates. (**a**) Non-social behaviour as measured by the duration and frequency of rearing up against the wall of the home-cage (frequency: *t*-test, t_24_ = 3.09, p = 0.01). (**b**) An inverse association between non-social behaviour and connectivity between retrosplenial and cingulate cortices (wall rearing, frequency: r = −0.45, p = 0.02, n = 26, *R*^2^ = 0.21). * p < 0.05, ** p < 0.01.

**Figure S7.**
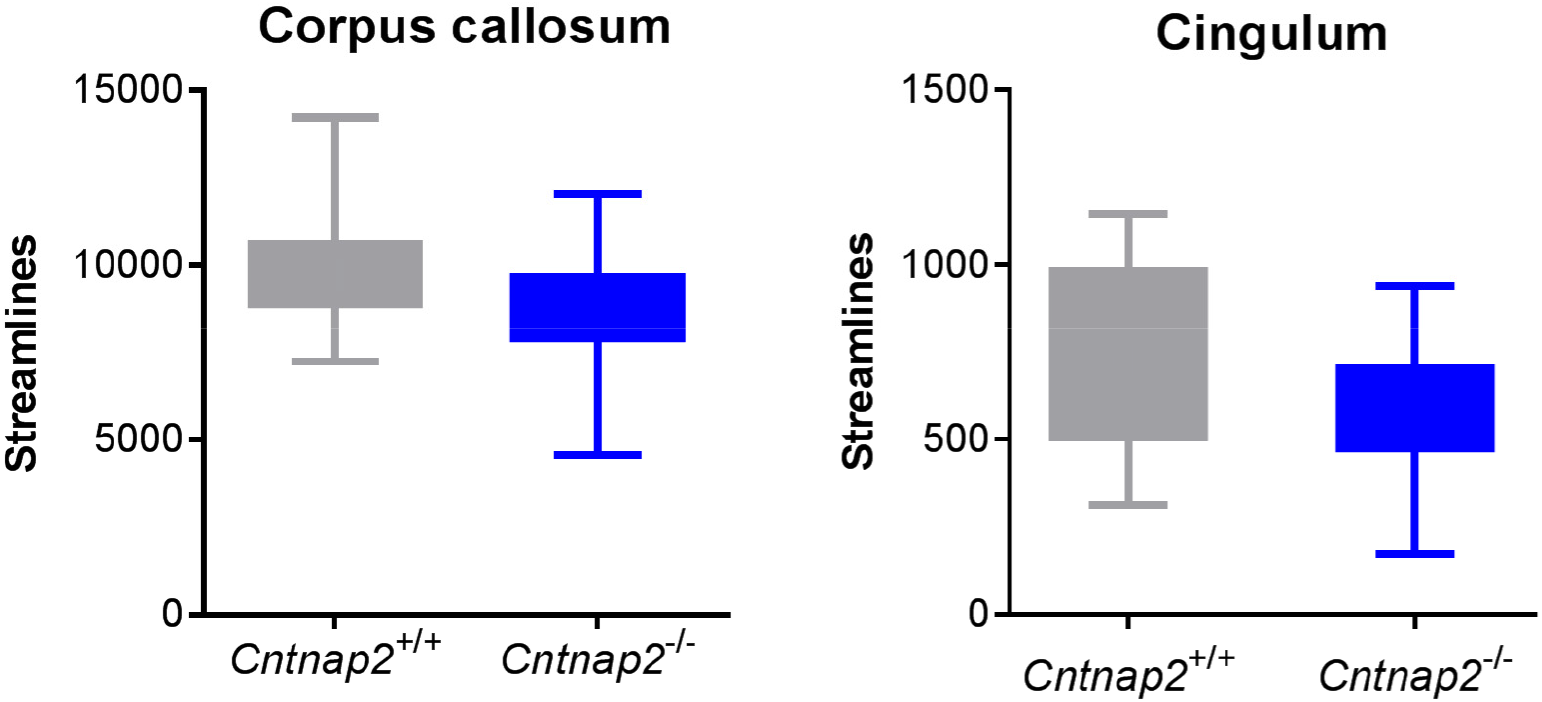
White-matter tractography-based streamline counts. Numbers of streamlines in corpus callosum and cingulum showed no significant differences between the *Cntnap2^−/−^* and control littermates (cingulum: *t*-test, t_21_ = 1.25, p = 0.23; corpus callosum: *t*-test, t_21_ = 1.21, p = 0.24).

**Figure S8.**
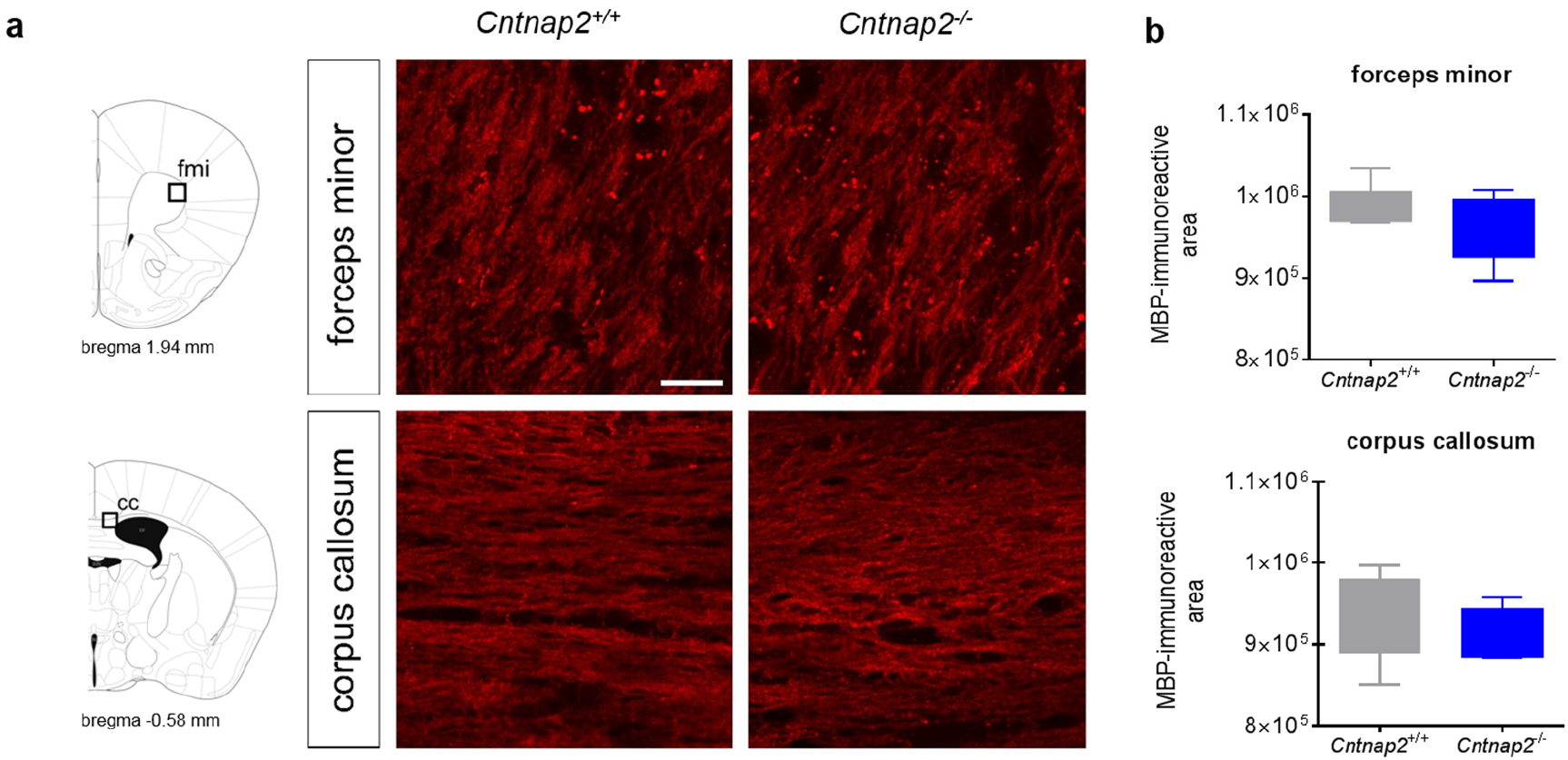
Histological and immunohistochemical analysis of white matter. (**a**) Representative images of anterior callosal regions characterized by parallel or transversal fibre extension with respect to the imaging plane (corpus callosum and forceps minor of the corpus callosum, respectively). No apparent difference in fibre organization or MBP stained regions was observed between genotypes. (**b**) MBP-immunoreactive area averaged from three random image fields per region and animal (*n* = 5, each group; corpus callosum: *t*-test, t_8_ = 0.84, p = 0.42; forceps minor of the corpus callosum: *t*-test, t_8_ = 1.06, p = 0.32).

